# HydDB: A web tool for hydrogenase classification and analysis

**DOI:** 10.1101/061994

**Authors:** Søndergaard Dan, Pedersen Christian N. S., Greening Chris

## Abstract

H2 metabolism is proposed to be the most ancient and diverse mechanism of energy-conservation. The metalloenzymes mediating this metabolism, hydrogenases, are encoded by over 60 microbial phyla and are present in all major ecosystems. We developed a classification system and web tool, HydDB, for the structural and functional analysis of these enzymes. We show that hydrogenase function can be predicted by primary sequence alone using an expanded classification scheme (comprising 29 [NiFe], 8 [FeFe], and 1 [Fe] hydrogenase classes) that defines 11 new classes with distinct biological functions. Using this scheme, we built a web tool that rapidly and reliably classifies hydrogenase primary sequences using a combination of *k*-nearest neighbors’ algorithms and CDD referencing. Demonstrating its capacity, the tool reliably predicted hydrogenase content and function in 12 newly-sequenced bacteria, archaea, and eukaryotes. HydDB provides the capacity to browse the amino acid sequences of 3248 annotated hydrogenase catalytic subunits and also contains a detailed repository of physiological, biochemical, and structural information about the 38 hydrogenase classes defined here. The database and classifier are freely and publicly available at http://services.birc.au.dk/hyddb/

## Introduction

Microorganisms conserve energy by metabolizing H2. Oxidation of this high-energy fuel yields electrons that can be used for respiration and carbon-fixation. This diffusible gas is also produced in diverse fermentation and anaerobic respiratory processes ^1^. H_2_ metabolism contributes to the growth and survival of microorganisms across the three domains of life: chemotrophs and phototrophs, lithotrophs and heterotrophs, aerobes and anaerobes, mesophiles and extremophiles alike ^1,2^. On the ecosystem scale, H_2_ supports microbial communities in most terrestrial, aquatic, and host-associated ecosystems ^1,3^. It is also proposed that H_2_ was the primordial electron donor ^4,5^. In biological systems, metalloenzymes known as hydrogenases are responsible for oxidizing and evolving H_2_ ^1,6^. Our recent survey showed there is a far greater number and diversity of hydrogenases than previously thought ^2^. It is predicted that over 55 microbial phyla and over a third of all microorganisms harbor hydrogenases ^2,7^. Better understanding H_2_ metabolism and the enzymes that mediate it also has wider implications, particularly in relation to human health and disease ^3,8^, biogeochemical cycling ^9^, and renewable energy ^10,11^.

There are three types of hydrogenase, the [NiFe], [FeFe], and [Fe] hydrogenases, that are distinguished by their metal composition. Whereas the [Fe]-hydrogenases are a small methanogenic-specific family ^12^, the [NiFe] and [FeFe] classes are widely distributed and functionally diverse. They can be classified through a hierarchical system into different groups and subgroups/subtypes with distinct biochemical features (e.g. directionality, affinity, redox partners, and localization) and physiological roles (i.e. respiration, fermentation, bifurcation, sensing) ^1,6^. It is necessary to define the subgroup or subtype of the hydrogenase to predict hydrogenase function. For example, while Group 2a and 2b [NiFe]-hydrogenases share > 35% sequence identity, they have distinct roles as respiratory uptake hydrogenases and H_2_ sensors respectively ^13,14^. Likewise, discrimination between Group A1 and Group A3 [FeFe]-hydrogenases is necessary to distinguish fermentative and bifurcating enzymes ^2,15^. Building on previous work ^16,17^, recently created a comprehensive hydrogenase classification scheme predictive of biological function ^2^. This scheme was primarily based on the topology of phylogenetic trees built from the amino acid sequences of hydrogenase catalytic subunits/domains. It also factored in genetic organization, metal-binding motifs, and functional information. This analysis identified 22 subgroups (within four groups) of [NiFe]-hydrogenases and six subtypes (within three groups) of [FeFe]-hydrogenases, each proposed to have unique physiological roles and contexts ^2^

In this work, we build on these findings to develop the first web database for the classification and analysis of hydrogenases. We developed an expanded classification scheme that captures the full sequence diversity of hydrogenase enzymes and predicts their biological function. Using this information, we developed a classification tool based on the *k*-nearest neighbors’ (*k*-NN) method. HydDB is a user-friendly, high-throughput, and functionally-predictive tool for hydrogenase classification that operates with precision exceeding 99.8%.

## Results and Discussion

### A sequence-based classification scheme for hydrogenases

We initially developed a classification scheme to enable prediction of hydrogenase function by primary sequence alone. To do this, we visualized the relationships between all hydrogenases in sequence similarity networks (SSN) ^18^, in which nodes represent individual proteins and the distances between them reflect BLAST *E*-values. As reflected by our analysis of other protein superfamilies ^19,20^, SSNs allow robust inference of sequence-structure-function relationships for large datasets without the problems associated with phylogenetic trees (e.g. long-branch attraction).Consistent with previous phylogenetic analyses ^2,16,17^, this analysis showed the hydrogenase sequences clustered into eight major groups (Groups 1 to 4 [NiFe]-hydrogenases, Groups A to C [FeFe]-hydrogenases, [Fe]-hydrogenases), six of which separate into multiple functionally-distinct subgroups or subtypes at narrower log*E* filters **(Figure 1; Figure S1)**. The SSNs demonstrated that all [NiFe]-hydrogenase subgroups defined through phylogenetic trees in our previous work ^2^ separated into distinct clusters, which is consistent with our evolutionary model that such hydrogenases diverged from a common ancestor to adopt multiple distinct functions ^2^. The only exception were the Group A [FeFe]-hydrogenases, which as previously-reported ^2,17^, cannot be classified by sequence alone as they have principally diversified through changes in domain architecture and quaternary structure. It remains necessary to analyze the organization of the genes encoding these enzymes to determine their specific function, e.g. whether they serve fermentative or electron-bifurcating roles.

**Figure. 1.** Sequence similarity network of hydrogenase sequences. Nodes represent individual proteins and the edges show the BLAST *E*-values between them at the log*E* filter defined at the bottom-left of each panel. The sequences are colored by class as defined in the legends. **Figure S1** shows the further delineation of the encircled [NiFe] hydrogenase classes.

The SSN analysis revealed that several branches that clustered together on the phylogenetic tree analysis ^2^ in fact separate into several well-resolved subclades **(Figure 1)**. We determined whether this was significant by analyzing the taxonomic distribution, genetic organization, metal-binding sites, and reported biochemical or functional characteristics of the differentiated subclades. On this basis, we concluded that 11 of the new subclades identified are likely to have unique physiological roles. We therefore refine and expand the hydrogenase classification to reflect the hydrogenases are more diverse in both primary sequence and predicted function than accounted for by even the latest classification scheme ^2^. The new scheme comprises 38 hydrogenase classes, namely 29 [NiFe]-hydrogenase subclasses, 8 [FeFe]-hydrogenase subtypes, and the monophyletic [Fe]-hydrogenases (**Table 1**).

**Table 1.** Expanded classification scheme for hydrogenase enzymes. The majority of the classes were defined in previous work ^2,16,17,46^. The [NiFe] Group 1i, 1j, 1j, 2e,4d, 4g, 4h, and 4i enzymes and [FeFe] Groups C1, C2, and C3 enzymes were defined in this work based on their separation into distinct clusters in the SSN analysis **(Figure 1)**. HydDB contains detailed information on each of these classes, including their taxonomic distribution, genetic organization, biochemistry, and structures, as well a list of primary references.

| **[NiFe] Group 1: Respiratory H<sub>2</sub>-uptake [NiFe]-hydrogenases** |  |  |  |
| --- | --- | --- | --- |
| **1a** | Periplasmic | Electron input for sulfate, metal, and organohalide respiration. [NiFeSe] variants. | 2 |
| **1b** | Prototypical | Electron input for sulfate, fumarate, metal, and nitrate respiration. | 2 |
| **1c** | Hyb-type | Electron input for fumarate, nitrate, and sulfate respiration. Physiologically reversible. | 2 |
| **1d** | Oxygen-tolerant | Electron input for aerobic respiration and oxygen-tolerant anaerobic respiration. | 2 |
| **1e** | Isp-type | Electron input primarily for sulfur respiration. Physiologically reversible. | 2 |
| **1f** | Oxygen-protecting | Unresolved role. May liberate electrons to reduce reactive oxygen species. | 2 |
| **1g** | Crenarchaeota-type | Electron input primarily for sulfur respiration. | 2 |
| **1h** | Actinobacteria-type | Electron input for aerobic respiration. Scavenges electrons from atmospheric H<sub>2</sub>. | 2,46 |
| **1i** | Coriobacteria-type (putative) | Undetermined role. May liberate electrons for anaerobic respiration. | This work |
| **1j** | Archaeoglobi-type | Electron input for sulfate respiration <sup>21</sup>. | This work |
| **1k** | Methanophenazine-reducing | Electron input for methanogenic heterodisulfide respiration <sup>22</sup>. | This work |
| **[NiFe] Group 2: Alternative and sensory uptake [NiFe]-hydrogenases** |  |  |  |
| **2a** | Cyanobacteria-type | Electron input for aerobic respiration. Recycles H<sub>2</sub> produced by other cellular processes. | 16 |
| **2b** | Histidine kinase-linked | H<sub>2</sub> sensing. Activates two-component system controlling hydrogenase expression. | 16 |
| **2c** | Diguanylate cyclase-linked (putative) | Undetermined role. May sense H<sub>2</sub> and regulate processes through cyclic di-GMP production. | 2 |
| **2d** | Aquificae-type | Unresolved role. May generate reductant for carbon fixation or have a regulatory role. | 2 |
| **2e** | Metallosphaera-type (putative) | Undetermined role. May liberate electrons primarily for aerobic respiration <sup>26</sup>. | This work |
| **[NiFe] Group 3: Cofactor-coupled bidirectional [NiFe]-hydrogenases** |  |  |  |
| **3a** | F<sub>420</sub>-coupled | Couples oxidation of H<sub>2</sub> to reduction of F<sub>420</sub> during methanogenesis. Physiologically reversible. [NiFeSe] variants. | 16 |
| **3b** | NADP-coupled | Couples oxidation of NADPH to evolution of H<sub>2</sub>. Physiologically reversible. May have sulfhydrogenase activity. | 16 |
| **3c** | Heterodisulfide reductase-linked | Bifurcates electrons from H<sub>2</sub> to heterodisulfide and Fd<sub>ox</sub> in methanogens. [NiFeSe] variants. | 16 |
| **3d** | NAD-coupled | Interconverts electrons between H<sub>2</sub> and NAD depending on cellular redox state. | 16 |
| **[NiFe] Group 4: Respiratory H<sub>2</sub>-evolving [NiFe]-hydrogenases** |  |  |  |
| **4a** | Formate hydrogenlyase | Couples formate oxidation to fermentative H<sub>2</sub> evolution. May be H<sup>+</sup>-translocating. | 2 |
| **4b** | Formate-respiring | Respires formate or carbon monoxide using H<sup>+</sup> as electron acceptor. Na<sup>+</sup>-translocating via Mrp <sup>23</sup>. | This work |
| **4c** | Carbon monoxide-respiring | Respires carbon monoxide using H<sup>+</sup> as electron acceptor. H<sup>+</sup>-translocating. | 2 |
| **4d** | Ferredoxin-coupled, Mrp-linked | Couples Fd<sub>red</sub> oxidation to H<sup>+</sup> reduction. Na<sup>+</sup>-translocating via Mrp complex <sup>24</sup>. | This work |
| **4e** | Ferredoxin-coupled, Ech-type | Couples Fd<sub>red</sub> oxidation to H<sup>+</sup> reduction. Physiologically reversible via H<sup>+</sup>/Na<sup>+</sup> translocation. | 2 |
| **4f** | Formate-coupled (putative) | Undetermined role. May couple formate oxidation to H<sub>2</sub> evolution and H<sup>+</sup> translocation. | 2 |
| **4g** | Ferredoxin-coupled (putative) | Undetermined role. May couple Fd<sub>red</sub> oxidation to proton reduction and H<sup>+</sup>/Na<sup>+</sup> translocation. | This work |
| **4h** | Ferredoxin-coupled, Eha-type | Couples Fd<sub>red</sub> oxidation to H<sup>+</sup> reduction in anaplerotic processes. H<sup>+</sup>/Na<sup>+</sup>-translocating <sup>25</sup>. | This work |
| **4i** | Ferredoxin-coupled, Ehb-type | Couples Fd<sub>red</sub> oxidation to H<sup>+</sup> reduction in anabolic processes. H<sup>+</sup>/Na<sup>+</sup>-translocating <sup>25</sup>. | This work |
| **[FeFe] Hydrogenases** |  |  |  |
| **A1** | Prototypical | Couples ferredoxin oxidation to fermentative or photobiological H<sub>2</sub> evolution. | 2,17 |
| **A2** | Glutamate synthase-linked (putative) | Undetermined role. May couple H<sub>2</sub> oxidation to NAD reduction, generating reductant for glutamate synthase. | 2,17 |
| **A3** | Bifurcating | Reversibly bifurcates electrons from H<sub>2</sub> to NAD and Fd<sub>ox</sub> in anaerobic bacteria. | 2,17 |
| **A4** | Formate dehydrogenase-linked | Couples formate oxidation to H<sub>2</sub> evolution. Some bifurcate electrons from H<sub>2</sub> to ferredoxin and NADP. | 2,17 |
| **B** | Colonic-type (putative) | Undetermined role. May couple Fd<sub>red</sub> oxidation to fermentative H<sub>2</sub> evolution. | 17 |
| **C1** | Histidine kinase-linked (putative) | Undetermined role. May sense H<sub>2</sub> and regulate processes via histidine kinases <sup>2</sup>. | This work |
| **C2** | Chemotactic (putative) | Undetermined role. May sense H<sub>2</sub> and regulate processes via methyl-accepting chemotaxis proteins <sup>2</sup>. | This work |
| **C3** | Phosphatase-linked (putative) | Undetermined role. May sense H<sub>2</sub> and regulate processes via serine/threonine phosphatases <sup>2</sup>. | This work |
| **[Fe] Hydrogenases** |  |  |  |
| **All** | Methenyl-H<sub>4</sub>MPT dehydrogenase | Reversibly couples H<sub>2</sub> oxidation to 5,10-methenyltetrahydromethanopterin reduction. | 16 |

Three lineages originally classified as Group 1a [NiFe]-hydrogenases were reclassified as new subgroups, namely those affiliated with Coriobacteria (Group 1i), Archaeoglobi (Group 1j), and Methanosarcinales (Group 1i). Cellular and molecular studies show these enzymes all support anaerobic respiration of H_2_, but differ in the membrane carriers (methanophenazine, menaquinone) and terminal electron acceptors (heterodisulfide, sulfate, nitrate) that they couple to ^21 22^. The previously-proposed 4b and 4d subgroups ^2^ were dissolved, as the SSN analysis confirmed they were highly polyphyletic. These sequences are reclassified here into five new subgroups: the formate- and carbon monoxide-respiring Mrp-linked complexes (Group 4b) ^23^, the ferredoxin-coupled Mrp-linked complexes (Group 4d) ^24^, the well-described methanogenic Eha (Group 4h) and Ehb (Group 4i) supercomplexes ^25^, and a more loosely clustered class of unknown function (Group 4g). Enzymes within these subgroups, with the exception of the uncharacterized 4g enzymes, sustain well-described specialist functions in the energetics of various archaea ^23–25^. Three crenarchaeotal hydrogenases were also classified as their own family (Group 2e); these enzymes enable certain crenarchaeotes to grow aerobically on O_2_ ^26,27^ and hence may represent a unique lineage of aerobic uptake hydrogenases currently underrepresented in genome databases. The Group C [FeFe]-hydrogenases were also separated into three main subtypes given they separate into distinct clusters even at relatively broad log*E* values **(Figure 1)**; these subtypes are each transcribed with different regulatory elements and are likely to have distinct regulatory roles ^2,17,28^ **(Table 1)**.

### HydDB reliably predicts hydrogenase class using the *k*-NN method and CDD referencing

Using this information, we built a web tool to classify hydrogenases. Hydrogenase classification is determined through a three-step process following input of the catalytic subunit sequence. Two checks are initially performed to confirm if the inputted sequence is likely to encode a hydrogenase catalytic subunit/domain. The Conserved Domain Database (CDD) ^29^ is referenced to confirm that the inputted sequence has a hydrogenase catalytic domain, i.e. “Complex1_49kDa superfamily” (cl21493) (for NiFe-hydrogenases), “Fe_hyd_lg_C superfamily” (cl14953) (for FeFe-hydrogenases), and “HMD” (pfam03201) (for Fe-hydrogenases). A homology check is also performed that computes the BLAST *E*-value between the inputted sequence and its closest homolog in HydDB. HydDB classifies any inputted sequence that lacks hydrogenase conserved domains or has low homology scores (*E*-value > 10^−5^) as a non-hydrogenase **(Table S1)**.

In the final step, the sequence is classified through the *k-NN* method that determines the most similar sequences listed in the HydDB reference database. To determine the optimal *k* for the dataset, we performed a 5-fold cross-validation for *k* = 1…10 and computed the precision for each *k*. The results are shown in **Figure 2**. The classifier predicted the classes of the 3248 hydrogenase sequences with 99.8% precision and high robustness when performing a 5-fold cross-validation (as described in the Methods section) for *k* = 4. The six sequences where there were discrepancies between the SSN and *k*-NN predictions are shown in **Table S2**. The classifier has also been trained to detect and exclude protein families that are homologous to hydrogenases but do not metabolize H2 (Nuo, Ehr, NARF, HmdII ^1,2^) using reference sequences of these proteins **(Table S1)**.

**Figure. 2.** Evaluating the *k*-NN classifier for *k* = 1…10. For each *k*, a 5-fold cross-validation was performed. The mean precision ± two standard deviations of the folds is shown in the figure (note the y-axis). *k* = 1 provides the most accurate classifier. However, *k* = 4 provides almost the same precision and is more robust to errors in the training set (reflected by the lower standard deviation). In general, the standard deviation is very small, indicating that the predictions are robust to changes in the training data.

Sequences of the [FeFe] Group A can be classified into functionally-distinct subtypes (A1, A2, A3, A4) based on genetic organization ^2^. The classifier can classify such hydrogenases if the protein sequence immediately downstream from the catalytic subunit sequence is provided. The classifier references the CDD to search for conserved domains in the downstream protein sequence. A sequence is classified as [FeFe] Group A2 if one of the domains “GltA”, “GltD”, “glutamate synthase small subunit” or “putative oxidoreductase”, but not “NuoF”, is found in the sequence. Sequences are classified as [FeFe] Group A3 if the domain “NuoF” is found and [FeFe] Group A4 if the domain “HycB” is present. If none of the domains are found, the sequence is classified as A1. These classification rules were determined by collecting 69 downstream protein sequences. The sequences were then submitted to the CDD and the domains which most often occurred in each subtype were extracted.

In addition to its precision, the classifier is superior to other approaches due to its usability. It is accessible as a free web service at http://services.birc.au.dk/hyddb/ HydDB allows the users to paste or upload sequences of hydrogenase catalytic subunit sequences in FASTA format and run the classification **(Figure S2)**. When analysis has completed, results are presented in a table that can be downloaded as a CSV file **(Figure S3)**. This provides an efficient and user-friendly way to classify hydrogenases, in contrast to the previous standard which requires visualization of phylogenetic trees derived from multiple sequence alignments ^30^.

### HydDB infers the physiological roles of H_2_ metabolism

As summarized in **Table 1**, hydrogenase class is strongly correlated with physiological role. As a result, the classifier is capable of predicting both the class and function of a sequenced hydrogenase. To demonstrate this capacity, we used HydDB to analyze the hydrogenases present in 12 newly-sequenced bacteria, archaea, and eukaryotes of major ecological significance. The classifier correctly classified all 24 hydrogenases identified in the sequenced genomes, as validated with SSNs **(Table 2)**. On the basis of these classifications, the physiological roles of H2 metabolism were predicted **(Table 2)**. For five of the organisms, these predictions are confirmed or supported by previously published data ^27,31–34^. Other predictions are in line with metabolic models derived from metagenome surveying ^35–37^. In some cases, the capacity for organisms to metabolize H2 was not tested or inferred in previous studies despite the presence of hydrogenases in the sequenced genomes ^32,38–40^.

**Table 2.** Predictive capacity of the HydDB. HydDB accurately determined hydrogenase content and predicted the physiological roles of H_2_ metabolism in 12 newly-sequenced archaeal and bacterial species.

| Organism | Phylum | Hydrogenase accession no. | HydDB classification | SSN classification | Predicted H<sub>2</sub> metabolism | Confirmed H<sub>2</sub> metabolism |
| --- | --- | --- | --- | --- | --- | --- |
| *Pyrinomonas methylaliphatogenes* | Acidobacteria | WP_041979300.1 | [NiFe] Group 1h | [NiFe] Group 1h | Persistence by aerobic respiration of atmospheric H<sub>2</sub> | Confirmed experimentally <sup>31</sup> |
| *Phaeodactylibacter xiamenensis* | Bacteroidetes | WP_044227713.1<br>WP_044216927.1<br>WP_044227053.1 | [NiFe] Group 1d<br>[NiFe] Group 2a<br>[NiFe] Group 3d | [NiFe] Group 1d<br>[NiFe] Group 2a<br>[NiFe] Group 3d | Chemolithoautotrophic growth by aerobic H<sub>2</sub> oxidation | Bacterium grows aerobically, but H<sub>2</sub> oxidation untested <sup>32</sup> |
| *Bathyarchaeota archaeon BA1* | Bathyarchaeota | KPV62434.1<br>KPV62673.1<br>KPV62298.1 | [NiFe] Group 3c<br>[NiFe] Group 3c<br>[NiFe] Group 4g | [NiFe] Group 3c<br>[NiFe] Group 3c<br>[NiFe] Group 4g | Couples Fd<sub>red</sub> oxidation to H<sub>2</sub> evolution in energy-conserving and bifurcating processes | Unconfirmed but consistent with metagenome-based models <sup>36</sup> |
| *Lenisia limosa* | Obozoa (Breviatea class) | LenisMan28 | [FeFe] Group A1 | [FeFe] Group A | Fermentative evolution of H<sub>2</sub> | Confirmed experimentally <sup>47</sup> |
| *Acidianus copahuensis* | Crenarchaeota | WP_048100721.1<br>WP_048100713.1<br>WP_048100378.1<br>WP_048100359.1 | [NiFe] Group 1g<br>[NiFe] Group 1g<br>[NiFe] Group 1h<br>[NiFe] Group 2e | [NiFe] Group 1g<br>[NiFe] Group 1g<br>[NiFe] Group 1h<br>[NiFe] Group 2e | Chemolithoautotrophic growth by H<sub>2</sub> oxidation using O<sub>2</sub> or S<sub>0</sub> as electron acceptors | Partially confirmed experimentally <sup>27</sup> |
| *Arcobacter* sp. E1/2/3 | Proteobacteria (Epsilon class) | Arc.peg.2312 | [NiFe] Group 1b | [NiFe] Group 1b | Chemolithoautotrophic growth by anaerobic H<sub>2</sub> oxidation | Confirmed experimentally <sup>47</sup> |
| *Methanoperedens nitroreducens* | Euryarchaeota (ANME) | WP_048088262.1<br>WP_048090768.1 | [NiFe] Group 3b<br>[NiFe] Group 3b | [NiFe] Group 3b<br>[NiFe] Group 3b | Secondary role for H<sub>2</sub> metabolism limited to fermentative evolution of H<sub>2</sub> | Unconfirmed but consistent with metagenome-based models <sup>35</sup> |
| *Kryptonium thompsoni* | Kryptonia | CUU03002.1<br>CUU06124.1 | [NiFe] Group 1d<br>[NiFe] Group 3b | [NiFe] Group 1d<br>[NiFe] Group 3b | Chemolithoautotrophic growth by aerobic H<sub>2</sub> oxidation, fermentative evolution of H<sub>2</sub>. | Untested, candidate phylum identified by metagenomics <sup>39</sup> |
| *Lokiarchaeum* sp. GC14_75 | Lokiarchaeota | KKK40681.1 | [NiFe] Group 3c | [NiFe] Group 3c | Bifurcates electrons between H<sub>2</sub>, heterodisulfide, and ferredoxin | Unconfirmed but consistent with metagenome-based models <sup>48</sup> |
| *Nitrospira moscoviensis* | Nitrospirae | WP_053379275.1 | [NiFe] Group 2a | [NiFe] Group 2a | Chemolithoautotrophic growth by aerobic H<sub>2</sub> oxidation | Confirmed experimentally <sup>33</sup> |
| *Bacterium* GW2011_GWE1_35_17 | Moranbacteria | KKQ46070.1<br>KKQ45273.1 | [NiFe] Group 1a<br>[NiFe] Group 3b | [NiFe] Group 1a<br>[NiFe] Group 3b | Chemolithoautotrophic growth by anaerobic H<sub>2</sub> oxidation, fermentative evolution of H<sub>2</sub>. | Unconfirmed but consistent with metagenome-based models <sup>37</sup> |
| *Bacterium* GW2011_GWA2_33_10 | Peregrinibacteria | KKP36897.1 | [FeFe] Group A3 | [FeFe] Group A | Bifurcates electrons between H<sub>2</sub>, NADH, and ferredoxin | Unconfirmed but consistent with metagenome-based models <sup>37</sup> |
| *Entotheonella* sp. TSY1 | Tectomicrobia | ETW97737.1<br>ETW94065.1 | [NiFe] Group 1h<br>[NiFe] Group 3b | [NiFe] Group 1h<br>[NiFe] Group 3b | Persistence by aerobic respiration of atmospheric H<sub>2</sub>, fermentative evolution of H<sub>2</sub> | Untested, candidate phylum identified by metagenomics <sup>40</sup> |

While HydDB serves as a reliable initial predictor of hydrogenase class and function, further analysis is recommended to verify predictions. Hydrogenase sequences only provide organisms with the genetic capacity to metabolise H_2_; their function is ultimately modulated by their expression and integration within the cell ^1,41^. In addition, some classifications are likely to be overgeneralized due to lack of functional and biochemical characterization of certain lineages and sublineages. For example, it is not clear if two distant members of the Group 1h [NiFe]-hydrogenases *(Robiginitalea biformata, Sulfolobus islandicus)* perform the same H_2_-scavenging functions as the core group ^9^. Likewise, it seems probable that the Group 3a [NiFe]-hydrogenases of Thermococci and Aquificae use a distinct electron donor to the main class ^42^. Prominent cautions are included in the enzyme pages in cases such as these. HydDB will be updated when literature is published that influences functional assignments.

### HydDB contains interfaces for hydrogenase browsing and analyzing

In addition to its classification function, HydDB is designed to be a definitive repository for hydrogenase retrieval and analysis. The database presently contains entries for 3248 hydrogenases, including their NCBI accession numbers, amino acid sequences, hydrogenase classes, taxonomic affiliations, and predicted behavior **(Figure S4)**. To enable easy exploration of the data set, the database also provides access to an interface for searching, filtering, and sorting the data, as well as the capacity to download the results in CSV or FASTA format. There are individual pages for the 38 hydrogenase classes defined here **(Table 1)**, including descriptions of their physiological role, genetic organization, taxonomic distribution, and biochemical features. This is supplemented with a compendium of structural information about the hydrogenases, which is integrated with the Protein Databank (PDB), as well as a library of over 500 literature references **(Figure S5)**.

## Conclusions

To summarize, HydDB is a definitive resource for hydrogenase classification and analysis. The classifier described here provides a reliable, efficient, and convenient tool for hydrogenase classification and functional prediction. HydDB also provides browsing tools for the rapid analysis and retrieval of hydrogenase sequences.Finally, the manually-curated repository of class descriptions, hydrogenase structures, and literature references provides a deep but accessible resource for understanding hydrogenases.

## Methods

### Sequence datasets

The database was constructed using the amino acid sequences of all curated non-redundant 3248 hydrogenase catalytic subunits represented in the NCBI RefSeq database in August 2014 ^2^ **(Dataset S1)**. In order to test the classification tool, additional sequences from newly-sequenced archaeal and bacterial phyla were retrieved from the Joint Genome Institute’s Integrated Microbial Genomes database ^43^.

### Sequence similarity networks

Sequence similarity networks (SSNs) ^18^ constructed using Cytoscape 4.1 ^44^ were used to visualize the distribution and diversity of the retrieved hydrogenase sequences. In this analysis, each node represents one of the 3248 hydrogenase sequences in the reference database **(Dataset S1)**. Each edge represents the sequence similarity between them as determined by *E*-values from all-vs-all BLAST analysis, with all self and duplicate edges removed. Three networks were constructed, namely for the [NiFe]-hydrogenase large subunit sequences **(Dataset S2)**, [FeFe]-hydrogenase catalytic domain sequences **(Dataset S3)**, and [Fe]-hydrogenase sequences **(Dataset S4)**. To control the degree of separation between nodes, logE cutoffs that were incrementally decreased from −5 to −200 until no major changes in clustering was observed. The log*E* cutoffs used for the final classifications are shown in **Figure 1** and **Figure S1**.

### Classification method

The *k*-NN method is a well-known machine learning method for classification ^45^. Given a set of data points *x_1_,x_2_, … x_N_* (e.g. sequences) with known labels *y_1_,y_2_*, …,*y_N_* (e.g. type annotations), the label of a point, *x,* is predicted by computing the distance from *x* to *x*_1_,*x*_2_, …*x_N_* and extracting the *k* labeled points closest to *x*, i.e. the neighbors. The predicted label is then determined by majority vote of the labels of the neighbors. The distance measure applied here is that of a BLAST search. Thus, the classifier corresponds to a homology search where the types of the top *k* results are considered. However, formulating the classification method as a machine learning problem allows the use of common evaluation methods to estimate the precision of the method and perform model selection. The classifier was evaluated using *k*-fold cross-validation. The dataset is first split in to *k* parts of equal size. *k* – 1 parts (the *training set)* are then used for training the classifier and the labels of the data points in the remaining part (the *test set)* are then predicted. This process, called a *fold,* is repeated *k* times. The predicted labels of each fold are then compared to the known labels and a precision can be computed.

## Acknowledgements

We thank A/Prof Colin J. Jackson, Dr Hafna Ahmed, Dr Andrew Warden, Dr Stephen Pearce, and the two anonymous reviewers for their helpful advice and comments regarding this manuscript. This work was supported by a PUMPkin Centre of Excellence PhD Scholarship awarded to DS and a CSIRO Office of the Chief Executive Postdoctoral Fellowship awarded to CG.

## Author Contributions

CG and DS designed experiments. DS and CG performed experiments. CG, DS,
and CNSP analyzed data. CNSP supervised students. CG and DS wrote the paper.

## Datasets

**Dataset S1.** Excel spreadsheet listing the sequence, taxonomy, and hydrogenase class of all 3248 hydrogenase catalytic subunit sequences listed in HydDB.

**Dataset S2.** Zip file containing the Cytoscape network for [NiFe]-hydrogenases.

**Dataset S3.** Zip file containing the Cytoscape network for [FeFe]-hydrogenases.

**Dataset S4.** Zip file containing the Cytoscape network for [Fe]-hydrogenases.

## Supporting Information

**Figure. S1.**
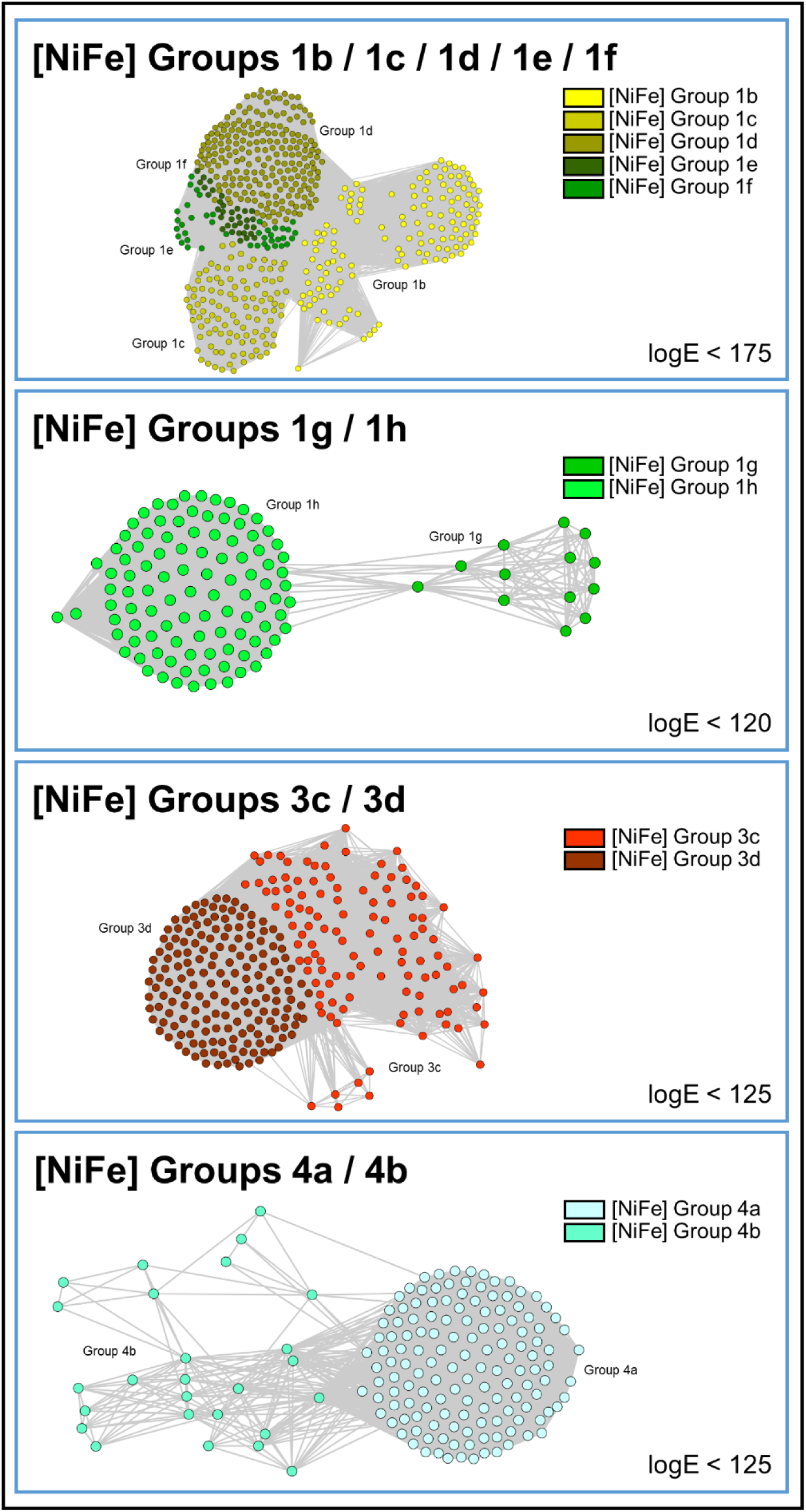
Sequence similarity networks showing the relationships between closely related subgroups of [NiFe]-hydrogenases as narrow log*E* filters.

**Figure. S2.**
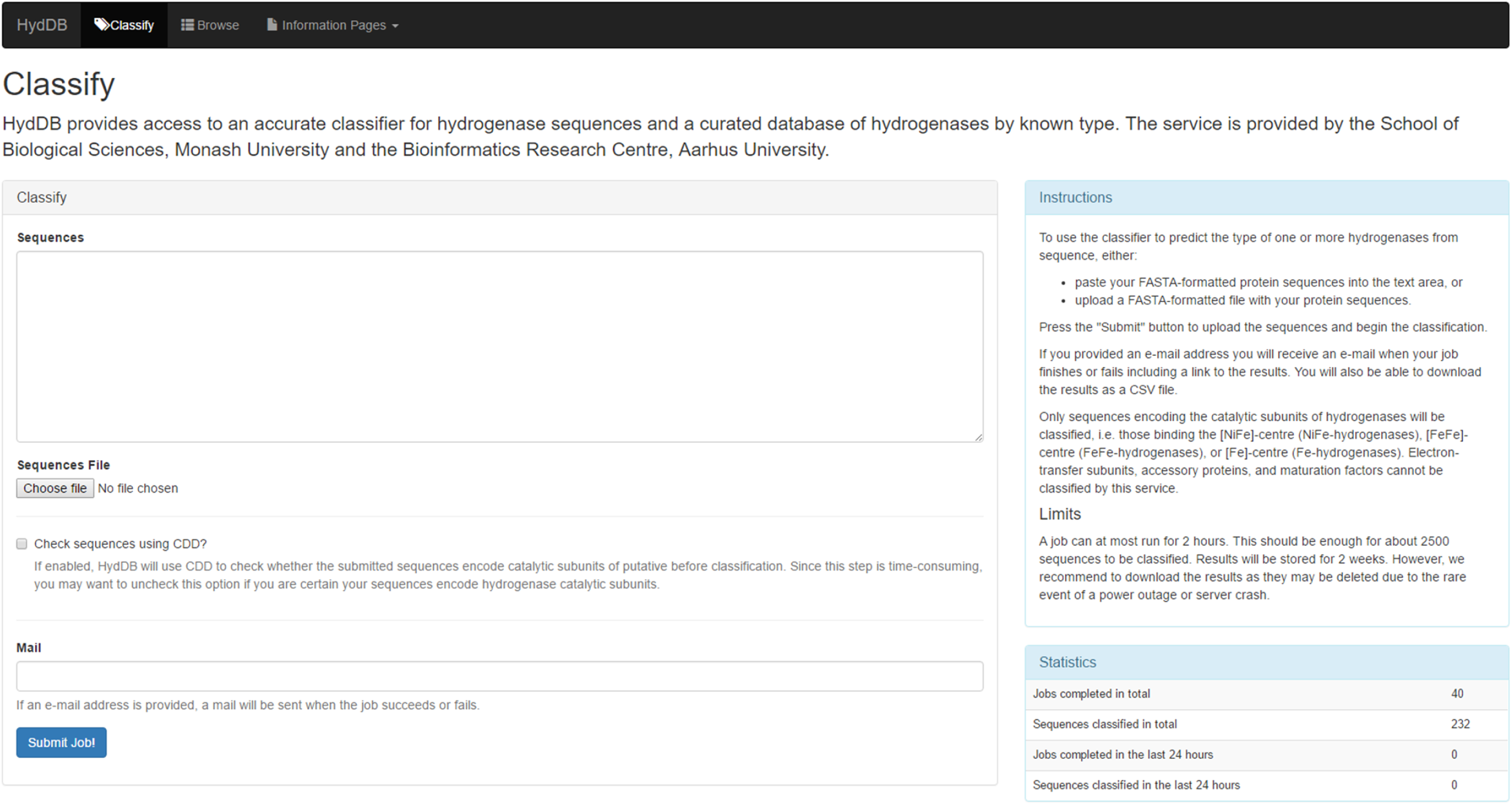
Screenshot showing interface of HydDB classification page.

**Figure. S3.**
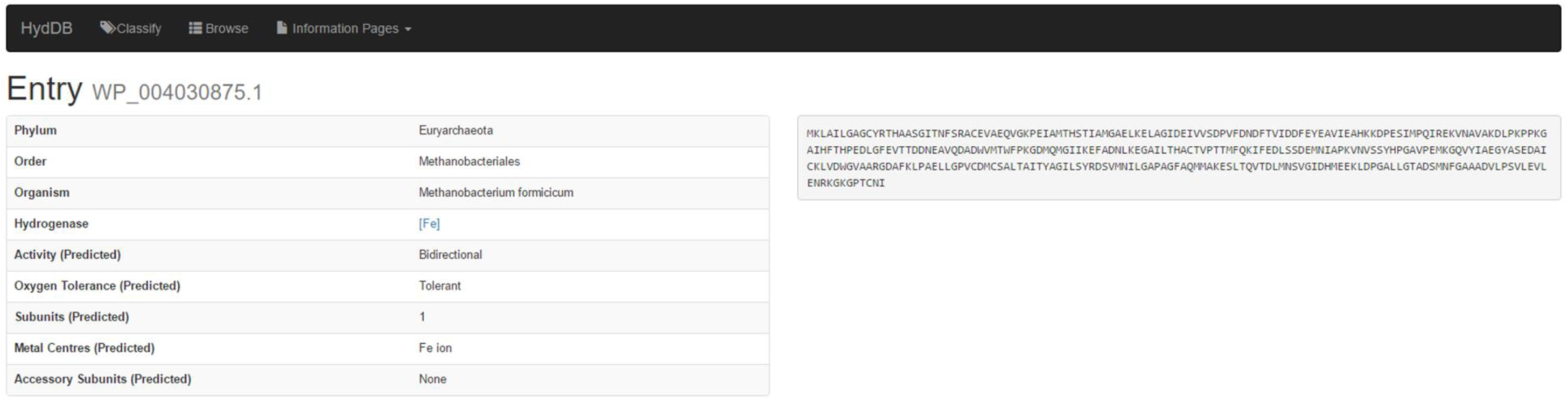
Screenshot showing the information provided in the data entry pages for 3248 individual hydrogenases in HydDB.

**Figure. S4.**
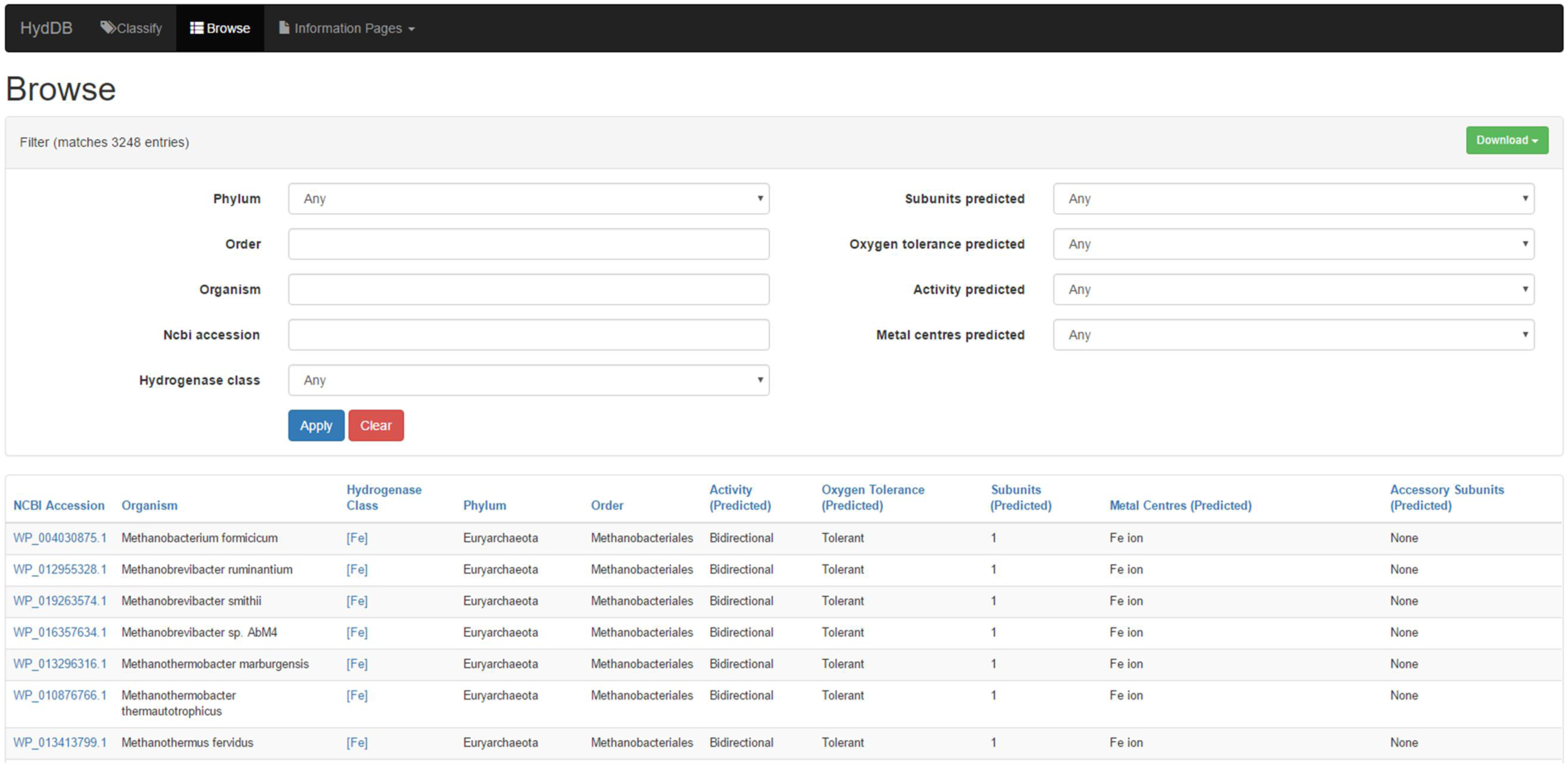
Screenshot showing the capacity for browsing hydrogenase data entries in HydDB.

**Figure. S5.**
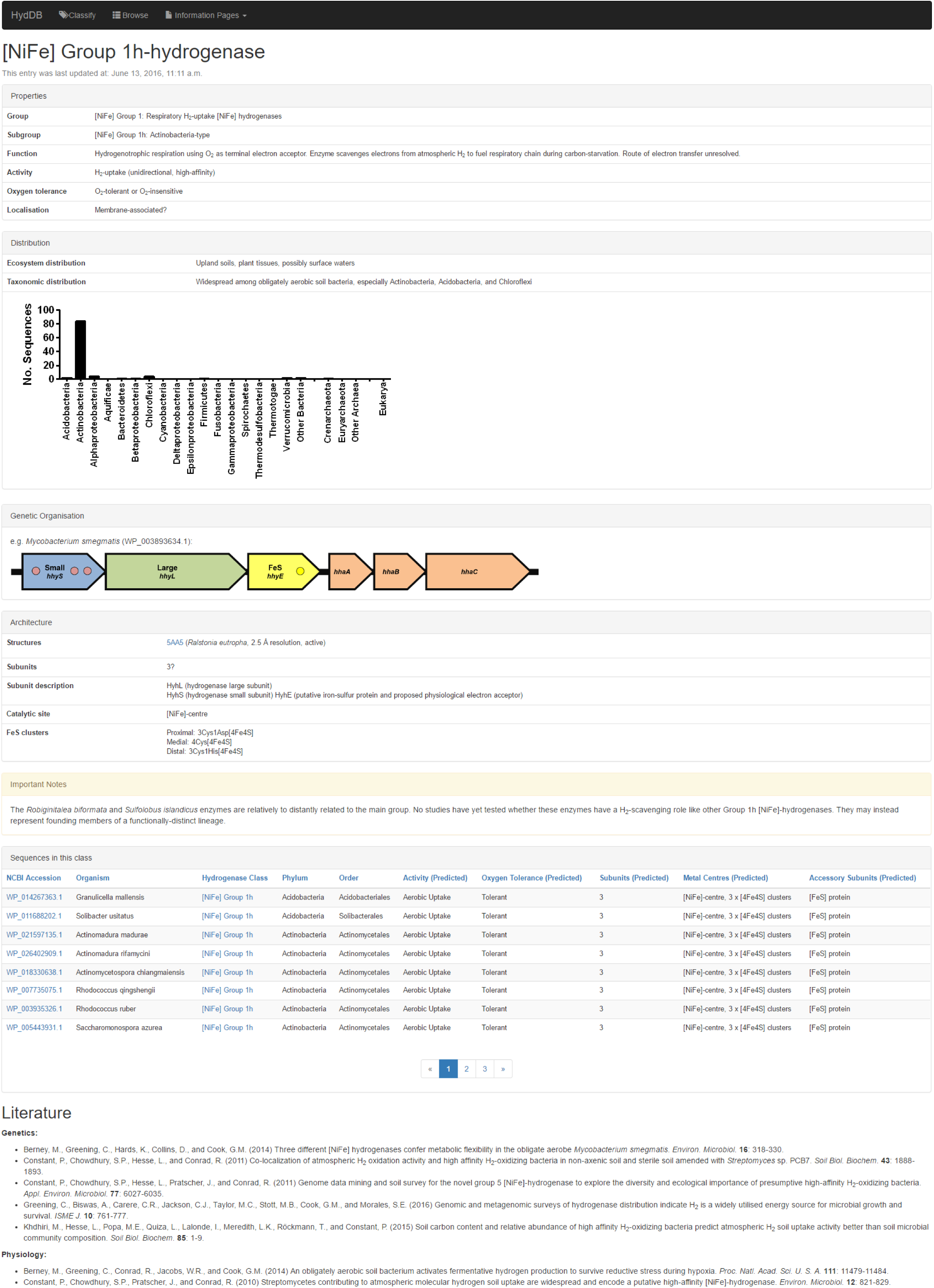
Screenshot showing the detailed content of the information pages about each hydrogenase class on HydDB. Equivalent information pages are available for all 38 hydrogenase classes defined in this work **(Table 1)**.

**Table S1.**
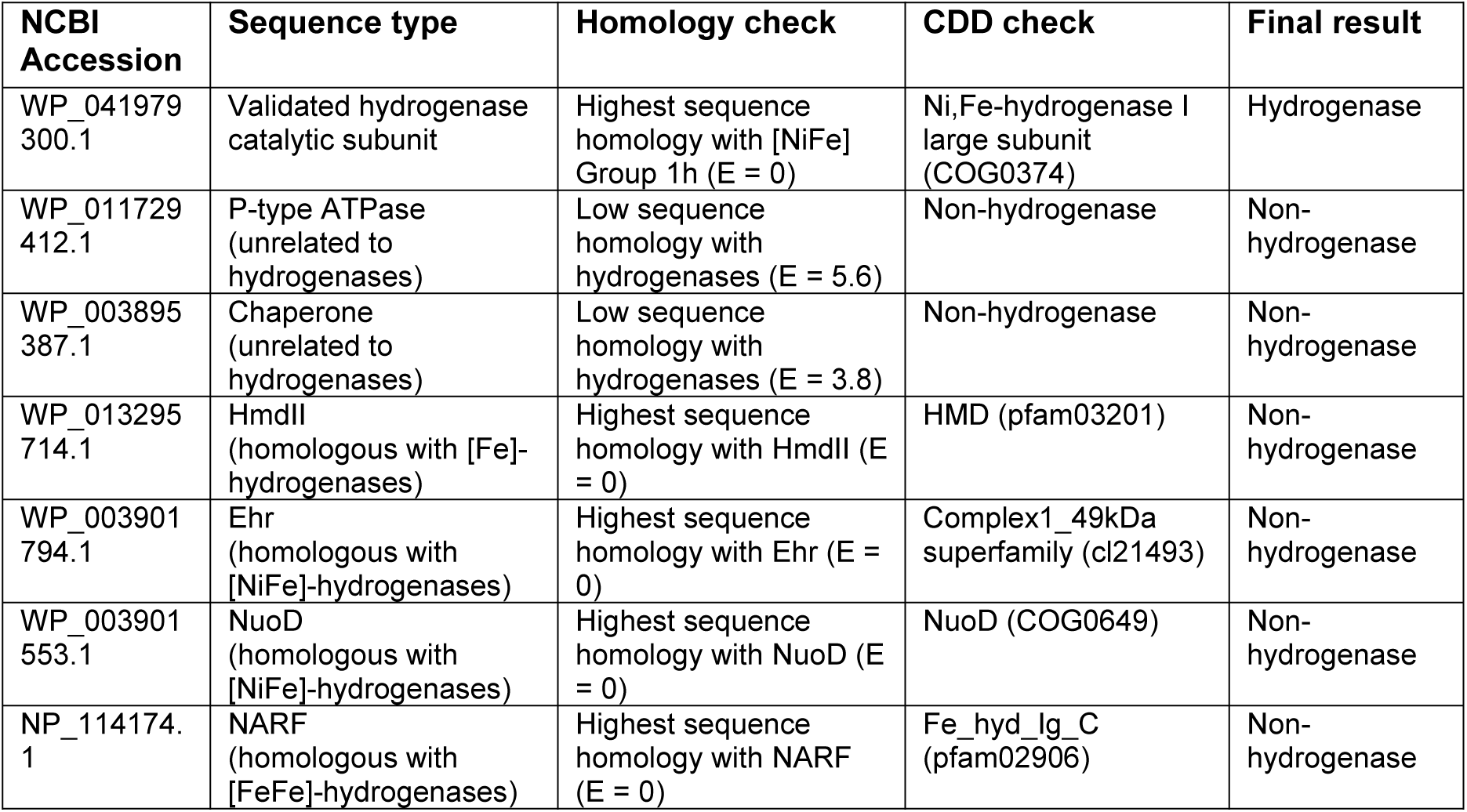
Validation that HydDB classifies only hydrogenase catalytic subunit sequences. HydDB excludes non-hydrogenase sequences through a combination of homology checks (sequences are only classified as hydrogenases if BLAST *E*-value of the closest hit in HydDB is less than 10^−5^) and CDD checks (sequences are only classified as hydrogenases if signature conserved domains are found). In addition, the classifier has been specifically trained to exclude four protein families that are homologous to hydrogenase catalytic subunits (HmdII, Her, NuoD, NARF) but lack hydrogenase activity.

| NCBI Accession | Sequence type | Homology check | CDD check | Final result |
| --- | --- | --- | --- | --- |
| WP_041979 300.1 | Validated hydrogenase catalytic subunit | Highest sequence homology with [NiFe] Group 1h ($E = 0$) | Ni,Fe-hydrogenase I large subunit (COG0374) | Hydrogenase |
| WP_011729 412.1 | P-type ATPase (unrelated to hydrogenases) | Low sequence homology with hydrogenases ($E = 5.6$) | Non-hydrogenase | Non-hydrogenase |
| WP_003895 387.1 | Chaperone (unrelated to hydrogenases) | Low sequence homology with hydrogenases ($E = 3.8$) | Non-hydrogenase | Non-hydrogenase |
| WP_013295 714.1 | HmdII (homologous with [Fe]-hydrogenases) | Highest sequence homology with HmdII ($E = 0$) | HMD (pfam03201) | Non-hydrogenase |
| WP_003901 794.1 | Ehr (homologous with [NiFe]-hydrogenases) | Highest sequence homology with Ehr ($E = 0$) | Complex1_49kDa superfamily (cl21493) | Non-hydrogenase |
| WP_003901 553.1 | NuoD (homologous with [NiFe]-hydrogenases) | Highest sequence homology with NuoD ($E = 0$) | NuoD (COG0649) | Non-hydrogenase |
| NP_114174.1 | NARF (homologous with [FeFe]-hydrogenases) | Highest sequence homology with NARF ($E = 0$) | Fe_hyd_Ig_C (pfam02906) | Non-hydrogenase |

**Table S2.** Hydrogenase sequences where there is disagreement between classification by SSN and *k*-NN methods. These sequences represent six out of the total 3248 sequences analyzed, i.e. 0.0018%.

| NCBI Accession | Organism | *k*-NN Classification | SSN Classification |
| --- | --- | --- | --- |
| WP_027414715.1 | Aneurinibacillus terranovensis | [NiFe] Group 1e | [NiFe] Group 1d |
| WP_027358538.1 | Desulforegula conservatrix | [NiFe] Group 3d | [NiFe] Group 3c |
| WP_012532312.1 | Geobacter bemidjiensis | [NiFe] Group 3d | [NiFe] Group 3c |
| WP_012469611.1 | Geobacter lovleyi | [NiFe] Group 3d | [NiFe] Group 3c |
| WP_004512544.1 | Geobacter metallireducens | [NiFe] Group 3d | [NiFe] Group 3c |
| WP_015839165.1 | Geobacter sp. M21 | [NiFe] Group 3d | [NiFe] Group 3c |

